# Poor Self-Reported Sleep is Associated with Prolonged White Matter T2 Relaxation in Psychotic Disorders

**DOI:** 10.1101/2024.07.03.601887

**Authors:** Haluk Umit Yesilkaya, Xi Chen, Lauren Watford, Emma McCoy, Ilgin Genc, Fei Du, Dost Ongur, Cagri Yuksel

## Abstract

**Background:** Schizophrenia (SZ) and bipolar disorder (BD) are characterized by white matter (WM) abnormalities, however, their relationship with illness presentation is not clear. Sleep disturbances are common in both disorders, and recent evidence suggests that sleep plays a critical role in WM physiology. Therefore, it is plausible that sleep disturbances are associated with impaired WM integrity in these disorders. To test this hypothesis, we examined the association of self-reported sleep disturbances with WM transverse (T2) relaxation times in patients with SZ spectrum disorders and BD with psychotic features.

**Methods:** 28 patients with psychosis (17 BD-I, with psychotic features and 11 SZ spectrum disorders) were included. Metabolite and water T2 relaxation times were measured in the anterior corona radiata at 4T. Sleep was evaluated using the Pittsburgh Sleep Quality Index.

**Results:** PSQI total score showed a moderate to strong positive correlation with water T2 (r = 0.64, p<0.001). Linear regressions showed that this association was specific to sleep disturbance but was not a byproduct of exacerbation in depressive, manic, or psychotic symptoms. In our exploratory analysis, sleep disturbance was correlated with free water percentage, suggesting that increased extracellular water may be a mechanism underlying the association of disturbed sleep and prolonged water T2 relaxation.

**Conclusion:** Our results highlight the connection between poor sleep and WM abnormalities in psychotic disorders. Future research using objective sleep measures and neuroimaging techniques suitable to probe free water is needed to further our insight into this relationship.

## INTRODUCTION

Schizophrenia (SZ) and Bipolar disorder (BD) overlap in genetic background, clinical presentation, and biological alterations (1, 2). In both disorders, an expanding body of literature indicates disruptions in white matter (WM) (3, 4), including in medication-free first-episode patients (5, 6) and unaffected relatives (7). The majority of evidence for WM pathology is derived from diffusion tensor imaging (DTI) studies, and the most commonly reported measure, fractional anisotropy (FA), does not provide information about the specific biological components affected (8). Nonetheless, additional lines of evidence, including other DTI measures, novel imaging techniques, and post-mortem and genetic studies suggest alterations in several aspects of WM microstructure, including axon, myelin, and extracellular water (3, 9-11). However, the link between WM abnormalities and illness presentation is not clear, as attempts to identify symptom correlates have largely been unfruitful.

Recent evidence suggests that sleep disturbances are associated with disrupted WM microstructure. In human neuroimaging studies, poor sleep was associated with altered FA and other diffusivity measures in the whole brain and specific WM tracts (12-20), and sleep deprivation was associated with widespread alterations in WM microstructure (21, 22). In addition, in primary insomnia disorder, FA was reduced in the internal capsule (23, 24), thalamus–pars triangularis tracts (25), and several regions, including the internal capsule, corona radiata, longitudinal fasciculus, and corpus callosum (26).

Sleep disturbances are highly prevalent in both SZ and BD, and are present even when patients are clinically stable or in the euthymic state (27, 28). However, despite the accumulating evidence indicating a role for sleep in WM physiology, little is known about the link between sleep disturbances and the WM disruptions observed in these disorders. To the best of our knowledge, there are no studies in schizophrenia that examined this relationship. In a recent study in individuals at ultra-high risk for psychosis, poor sleep was associated with lower FA in corpus callosum and both increased and decreased FA in the ventral brain regions (29). In BD, lower objective and self-reported sleep duration correlated with reduced FA and increased radial diffusivity (RD) in multiple white matter tracts (Benedetti et al., 2017). In contrast, in another study, poor sleep (reduced sleep duration, more sleep inertia) was associated with higher FA in several WM tracts (Verkooijen et al., 2017).

Compared to diffusion, which reflects the mean square displacement traveled by a molecule in unit time, T2 relaxation is determined by spin-spin interactions between the index molecule and its immediate neighbors, e.g. macromolecules. T2 relaxation time and diffusion measures reflect related but distinct aspects of the cellular microenvironment. Water T2 relaxation times can provide information on white matter macromolecule structure and fluid homeostasis while the metabolite relaxation times reflect the intra-axonal milieu. In a previous study, we observed increased water T2 relaxation time (T2R) as well as a reduced N-acetyl aspartate (NAA) T2R in chronic SZ compared to controls (30). Prolonged WM water T2R in SZ has also been reported in previous studies (31-33). We also observed that NAA T2R is significantly reduced in the chronic psychosis compared to first episode (FE) subjects, suggesting that apparent NAA concentration reductions reported in psychotic disorders may indeed reflect shortened T2R and not lower NAA tissue concentration (34). More recently, in a longitudinal study in FE psychosis, we observed a significant reduction of NAA in the second year of the follow-up compared to baseline, while the water T2 relaxation time showed a trend of increase (35).

Given this background, we hypothesized that sleep disturbances would be associated with white matter disruption in psychosis. To test this hypothesis, we examined the link between self-reported sleep quality and WM water and metabolite T2R in patients with psychotic disorders (SZ spectrum disorders and BD with psychotic features). Sleep supports neuronal integrity and neuroplasticity, as well as myelin physiology and brain fluid homeostasis. Therefore, we hypothesized that poor sleep quality would be associated with alterations in both metabolite and water T2R.

## MATERIALS AND METHODS

### Participants

This is a secondary analysis of data obtained in two studies, which acquired T2 data with the same protocols (30, 35). Patients were recruited from the inpatient and outpatient services at McLean Hospital. Men and women between 18-55 years old were included. Participants with any uncontrolled medical disorders, intellectual disability, neurological sequela, history of head trauma with loss of consciousness, and contraindication to MRI were excluded. The studies were approved by McLean Hospital and Mass General Brigham institutional review boards and all participants provided written informed consent. The study procedures adhered to the principles outlined in the Declaration of Helsinki. 28 patients (17 BD-I and 11 SZ spectrum disorders) provided information about sleep quality within 1 month of their scan (average interval 11.4±10.6 days) and were included in this study. The sample consisted of predominantly early-course patients, with 80.8% of the patients within the first three years of illness onset. Demographic and clinical information in this sample is displayed in **Table 1**.

**Table 1.** Clinical and demographic variables and T2R measures in the sample. Some of the data is presented as mean ± standard deviation. BD: Bipolar disorder; BMI: Body mass index; CPZ: chlorpromazine equivalent of antipsychotic dose; MADRS: Montgomery-Asberg Depression Rating Scale; PANSS: Positive and Negative Syndrome Scale; PSY-NOS: Psychotic disorder, not otherwise specified; SZA: Schizoaffective disorder; YMRS: Young Mania Rating Scale.

|  |  |  |
| --- | --- | --- |
| 316 |  |  |
| 317 | Age | 24.3±4.9 |
| 318 | Sex (%Females) | 35.7% |
| 319 |  |  |
| 320 | BMI | 24.9±4.6 |
| 321 |  |  |
| 322 | **Diagnosis and History** |  |
| 323 | SZA | 32.1% |
| 324 |  |  |
| 325 | Psy-NOS | 7.1% |
| 326 |  |  |
| 327 | BD-I | 60.7% |
| 328 |  |  |
| 329 | Duration of Illness (years) | 2.4±2 |
| 330 |  |  |
| 331 | **Medications** |  |
| 332 | Medication users | 85.7% |
| 333 |  |  |
| 334 | Antipsychotic users | 67.9% |
| 335 |  |  |
| 336 | CPZ | 225.9±226.9 |
| 337 |  |  |
| 338 | Lithium users | 42.9% |
| 339 |  |  |
| 340 | Other mood stabilizer users | 28.6% |
| 341 | **Symptom Severity** |  |
| 342 | YMRS | 6.5±8.1 |
| 343 |  |  |
| 344 | MADRS | 24.8±7.4 |
| 345 |  |  |
| 346 | PANSS total | 9.2±8.3 |
| 347 |  |  |
| 348 | PSQI total | 5.6±2.5 |
| 349 | **T2R** |  |
| 350 | Water | 64±3.4 |
| 351 |  |  |
| 352 | NAA | 255.7±23.8 |
| 353 |  |  |
| 354 | Cr | 148±12 |
| 355 |  |  |
| 356 | Cho | 175.5±22 |

### Clinical Assessments

Sleep was assessed using the Pittsburgh Sleep Quality Index (PSQI) (36), a self-report questionnaire that probes the sleep quality and disturbances over a 1-month period. The composite score, PSQI total score, was used in all analyses. A higher PSQI total score reflected poorer sleep. Diagnoses were ascertained using the Structured Clinical Interview for DSM-IV (SCID). In addition to PSQI, the severity of psychotic, manic, and depressive symptoms was assessed using the Positive and Negative Syndrome Scale (PANSS), the Young Mania Rating Scale (YMRS), and the Montgomery-Asberg Depression Rating Scale (MADRS). Antipsychotic load was calculated as the total chlorpromazine equivalent dose (CPZ) (37).

### MRI

T2 relaxation time measurements were conducted on a 4 T Varian full-body MR scanner (Unity/Inova; Varian NMR Instruments, CA, USA) using a 16-rung, single-tuned, volumetric birdcage coil. Global shimming was performed, followed by the acquisition of high-contrast T1-weighted sagittal images, which served to position the axial images and MRS voxels. A 1 × 3 × 3 cm^3^ single MRS voxel **(Figure 1**) was then placed on the corona radiata, centered at the level of the genu of the corpus callosum but lateral to it (i.e., does not include any callosal fibers). The voxel was placed in pure white matter with adjacent gray matter in the anterior and lateral directions used as anchors to ensure that the location was consistent across scans. SPM12 was used for tissue segmentation on the T1. The MRS voxel tissue percentages were calculated using AFNI and the voxel was consistently positioned in WM mostly (88 ± 4% of WM percentage). Localized shimming was performed to ensure water linewidths < 15 Hz.

**Figure 1.**
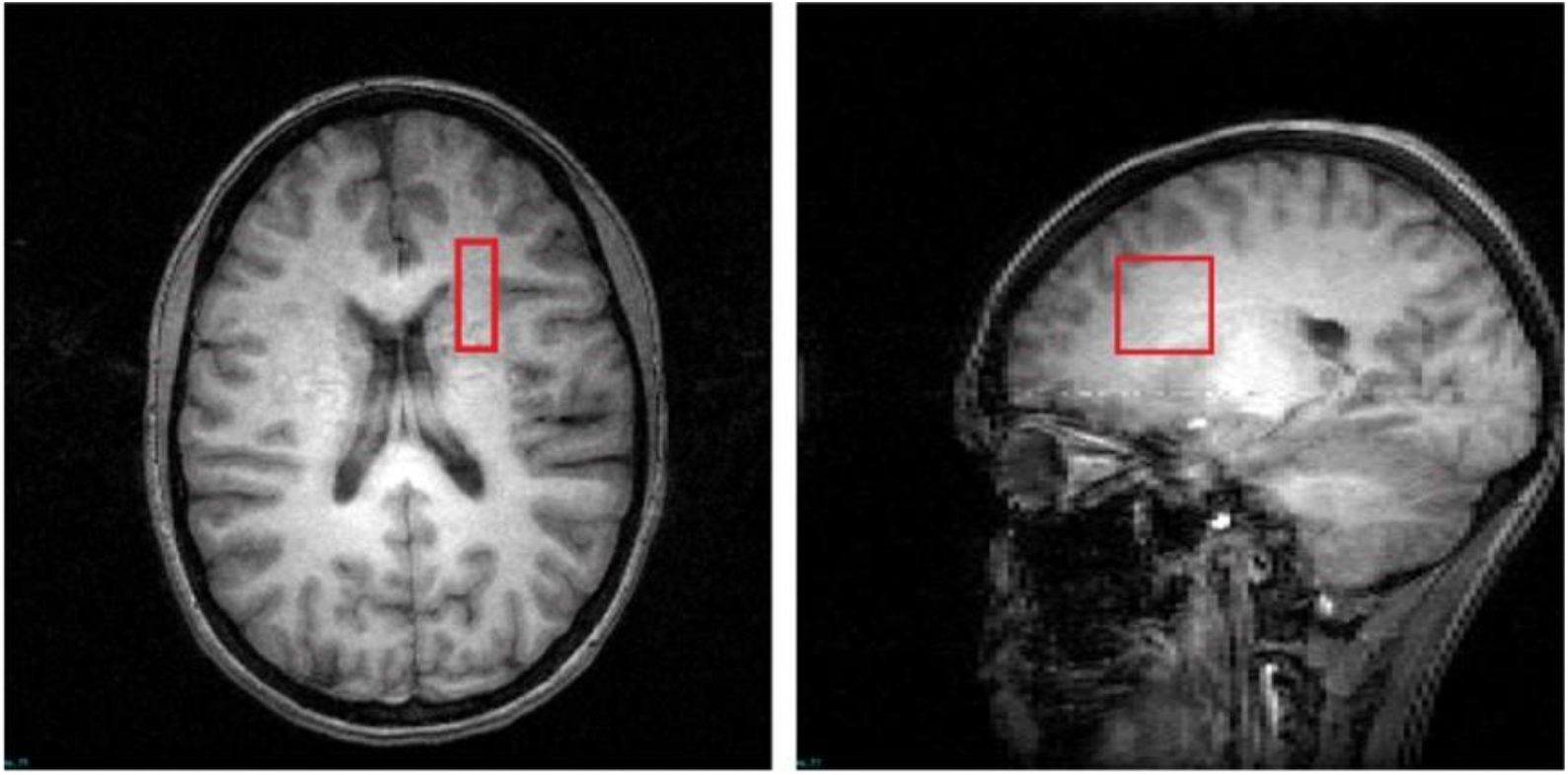
T1-weighted images in the transverse and sagittal planes depict the voxel placement.

Metabolite T2 spectra were obtained using a PRESS sequence modified with four varying TEs (30, 90, 120, and 200 ms) and TR = 3000 ms; 48 repetitions for metabolite and 8 repetitions for water T2 relaxation time measurements. A 3 ms sinc pulse with a bandwidth of 2000 Hz was used for excitation; two 6 ms Varian optc4 (Optimized Control Pulse for 4 zero sinc pulse) pulses with a bandwidth of 1050 Hz were used for refocusing.

We further performed a bi-exponential fitting for water T2 relaxometry to investigate the water compartments, as we found that the mono-exponential fitting may not fully account for water T2 decay **(Supplementary Figure S1)**. In the bi-exponential model of water T2 relaxometry, T2_fast (< 80 ms), T2_slow (> 120 ms), and the percentages of these two components were calculated. T2_fast, which reflects intra- and extra-cellular water relaxation, and the percentage of the slow component, which is considered as free water (FW%), were included in the exploratory analysis.

### Statistical Analysis

Statistical analyses were performed using IBM SPSS Statistics Version 26. Pearson or Spearman’s rank correlation was used to examine the correlations between sleep disturbance and T2 relaxation, depending on the distribution (normal vs. non-normal) and the type of (continuous vs. ordinal) data.

Linear regressions were used to test the associations of PSQI total score with neuroimaging measures of interest adjusted for demographic and clinical variables. Partial regression plots and a plot of studentized residuals against the predicted values indicated that assumptions of linear relationship were met. Histogram and P-P plot of standardized residuals showed a normal distribution. Variance inflation factors (VIF) indicated that no confounding multicollinearity was present.

All analyses were two-tailed. The significance level for hypothesis testing (α) was set at 0.05. For exploratory analyses, the Benjamini-Hochberg procedure was used to correct for multiple comparisons with the threshold for false discovery rate (FDR) at 0.05, and adjusted p values are presented.

## RESULTS

The average PSQI total score was 5.64, above the cut-off value of 5, which indicates “poor sleep” (36). PSQI total score was positively correlated with water T2R in the whole patient sample (r = 0.64, p = 10^−4^ × 2; **Figure 2**), and patients with BD-I and SZ spectrum disorders showed similar correlations (r = 0.63, p = 0.007 and r = 0.63, p = 0.038). There were no other significant correlations with the PSQI total score and any of the metabolite T2R values (all p>0.5).

**Figure 2.**
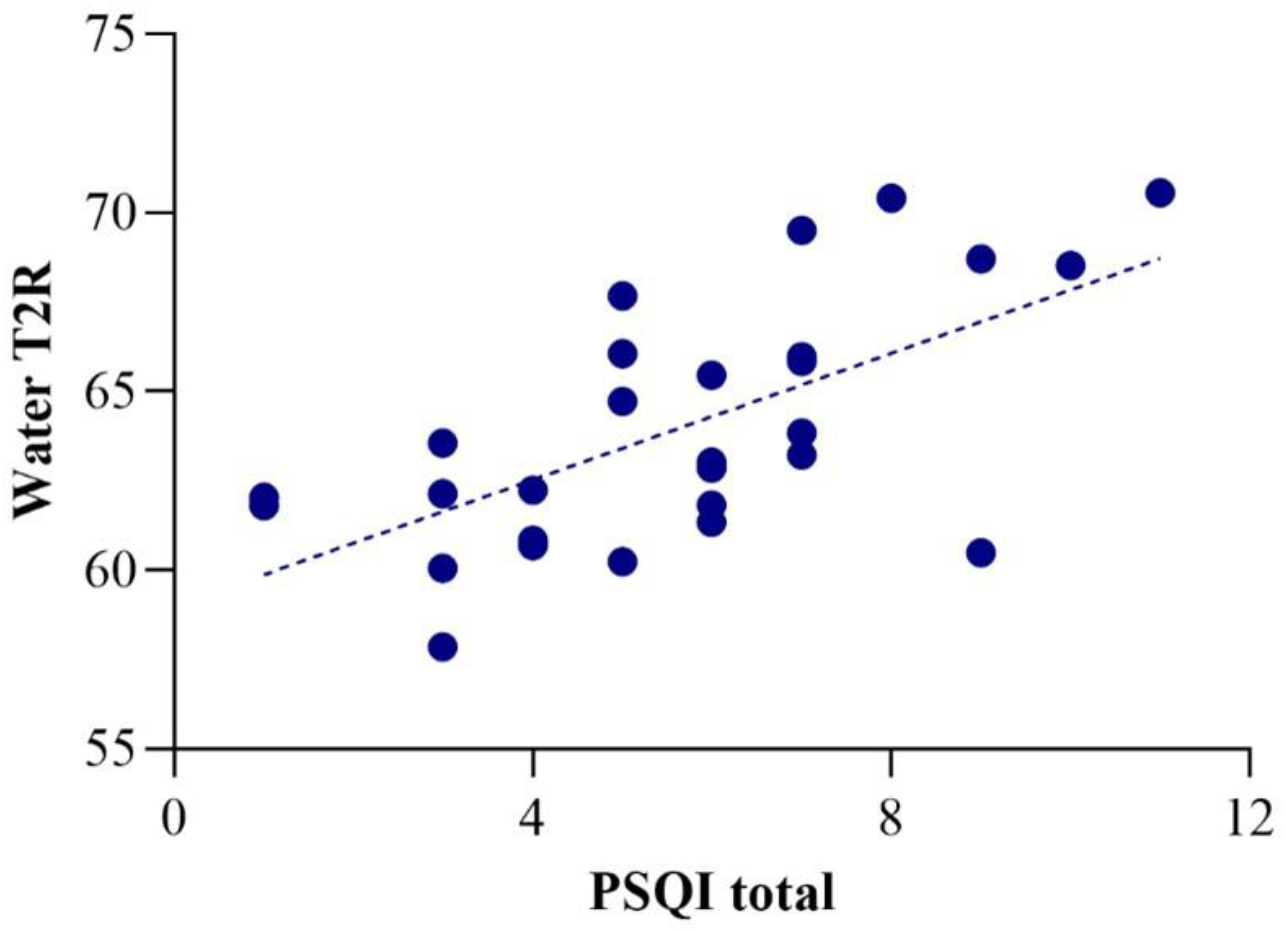
Correlation of water T2R with PSQI total score (r=0.64).

To see whether the association of sleep disturbance with water T2R was specific to this symptom dimension or simply a byproduct of increased overall symptom severity, we carried out separate linear regressions with PSQI total score included as a predictor along with symptom scores for mania (YMRS score), depression (MADRS score) or psychosis (PANSS total score). Sleep item scores were subtracted from YMRS and MADRS total scores. Age and sex were also included as additional covariates. These analyses showed that sleep disturbance remained a significant predictor of water-T2 (PSQI total and YMRS: β = 0.62, p = 0.002; PSQI total and MADRS: β = 0.62, p = 0.003; PSQI total and PANSS total: β = 0.64, p = 0.006), indicating that this association was independent of overall symptom severity.

We also found that the mono-exponential fitting quality was inversely correlated with PSQI in the patient group (spearman’s rho = -0.52, p = 0.006). The association between mono-exponential fitting quality and PSQI implies that additional compartments of water may affect the mono-exponential fitting quality and thus could be a biomarker of white matter integrity related to sleep quality. Therefore, we performed an exploratory analysis with bi-exponential fitting for water T2 decay and we found that PSQI total was positively correlated with FW% (spearman’s rho = 0.42, p = 0.032) **(Supplementary Figure S2)**, but not with T2_fast (r = 0.08, p = 0.666), suggesting that increased free water may be a potential mechanism underlying the association of poor sleep with prolonged water T2R.

## DISCUSSION

In this study, we examined the association of sleep quality with prefrontal WM water and metabolite T2R in patients with BD-I and SZ spectrum disorders. Supporting our hypothesis, we found that self-reported poor sleep was associated with prolonged water-T2R. Sleep quality was the only variable that was associated with water-T2R, and this association was independent of the severity of other manic, depressive, or psychotic symptoms.

Water-T2R reflects the interaction of water with nonaqueous molecules in its microenvironment and is prolonged in conditions where the frequency of these interactions is reduced due to the relative expansion of the water component (38). Prolonged WM water-T2R has been reported in SZ (30, 33, 39) with suggestions that such findings may arise from disruptions in myelin integrity, reduced axon size, or increased interstitial fluid. In line with the proposition that Water-T2R may reflect myelin abnormalities, the correlation of poor sleep with longer water-T2R in our sample is consistent with the expanding literature which indicates that sleep influences oligodendrocyte function, expression of numerous genes related to membrane metabolism and myelination, and sleep deprivation leads to myelin disruption (reviewed in 40). Furthermore, sleep is crucial for neuronal homeostasis and neuroplasticity (41-43) with poor sleep linked to reduced gray matter in the prefrontal cortex (PFC) (44-49), a region where the anterior corona radiata fibers project to. In addition, abnormal sleep and experimental sleep deprivation lead to impairments in executive functions which are mediated by the PFC (47, 50-52) and associated with corona radiata microstructure (53, 54). Consequently, it is plausible that sleep disruption-related neuronal alterations in the PFC are accompanied by reduced axon size in corona radiata, thereby manifesting as a relative expansion of the water component. However, the lack of correlation between sleep disturbance and N-acetylaspartate (NAA) T2R in our sample argues against this possibility, as a decrease in axon size would be expected to prolong the T2 relaxation for intracellular metabolites.

The positive correlation of FW% with PSQI total score suggests that increased FW due to poor sleep is a plausible mechanism, which would be consistent with a recent study (20). An increase in FW has been documented in the early course SZ patients (55-60) and in individuals at clinical high risk for psychosis (61), and has been shown to be inversely correlated with the duration of illness (60). Our sample consisted predominantly of early-course patients, which supports this possibility. Other than free water imaging based on DTI (62, 63), multi-exponential T2 relaxometry has been a potentially useful technique for characterizing the tissue water compartments (64, 65). The compartment with the shortest T2 (10 – 20 ms) is usually regarded as myelin water (66, 67) and it can hardly be detected with the current T2 spectroscopy protocol as our shortest TE is 30 ms. The main water compartment observed by the current study is with T2 = 40 – 80 ms and it is mainly contributed by intra- and extra-cellular water (68). The T2 values we obtained from the mono-exponential fitting mostly reflected the T2 decay of this compartment. Another water compartment observed by the bi-exponential fitting is with T2 > 120 ms and is usually regarded as free water (68, 69). It has a much lower fraction compared to intra- and extra-cellular water while it brings long tails to the T2 decay curves (70) and deviations from the mono-exponential fitting. It should be noted that the T2s of this compartment (T2_slow) in the current study are < 400 ms and thus are not contributed by CSF, which has a very long T2 from 800 - 3000 ms and minimal tissue percentages of the MRS voxel in the current study (0.1 ± 0.1%). Neuroinflammation has been proposed as a potential mechanism that leads to increased FW in SZ (60), and consistent with this hypothesis, previous investigations showed that increased peripheral levels of pro-inflammatory cytokines in SZ are associated with the expansion of the FW compartment (71-73). Notably, there is substantial evidence indicating that sleep disturbance and duration are linked to systemic inflammation (74, 75). Finally, recent evidence suggests that brain fluid dynamics are tightly linked to sleep-wake states, with increased cerebrospinal fluid (CSF) influx and significant augmentation of interstitial fluid observed during sleep (76). Although this burgeoning area of research has not been explored in SZ or BD, it is conceivable that abnormal CSF or glymphatic system dynamics due to poor sleep contribute to the observed increase in FW. Given the cross-sectional design of our study, we cannot speculate on the causal directionality of these potential mechanisms.

Our study had several limitations. First, given the constraints on experiment time, we have acquired signals of only 4 different TEs to measure T2. With limited data points, the exploratory bi-exponential analysis could be subject to inaccuracy and is not able to address the contribution from the ultra-short T2 components such as myelin. On the other hand, we acquired the full FID signal of each TE from a well-shimmed MRS voxel instead of just a few signal points using fast multi-echo imaging to achieve better signal reliability. The Carr-Purcell-Meiboom-Gill (CPMG) method used in multi-echo imaging is also sensitive to inhomogeneous B1 (RF) and B0 (static) fields (67, 69). Despite the limited number of TEs, our bi-exponential fitting showed significant improvement compared to mono-exponential **(Supplementary Figure S1)**. Nonetheless, given this limitation, the correlation of poor sleep with FW% should be treated as a preliminary finding offering potential guidance for future investigations exploring this relationship using suitable neuroimaging techniques. Second, because of the relatively small sample size and to avoid inflating the error rate, we could not take into account other potential factors that could affect WM T2R, such as body mass index. However, our previous studies (30, 35), from which we derived the current study’s data, did not find any association of WM water T2R with demographic or clinical factors in larger samples, except for sex, which is already controlled for in the regressions. Third, we did not have any objective sleep data, therefore, we could not explore the convergence with the self-reported sleep disturbances. Finally, while all the participants were psychotic, our sample consisted mostly of patients with affective symptoms (SZA and BD). Therefore, future studies should investigate if similar associations are present in “non-affective” psychosis.

In conclusion, our findings suggest that poor sleep quality is associated with WM abnormalities in patients with psychotic disorders. Increased free water, possibly due to neuroinflammation, is a plausible mechanism underlying this association. Future studies should include additional objective sleep measures and specialized neuroimaging techniques that probe the free water component in the WM.

## Acknowledgments

This project was supported by the National Institute of Mental Health K23MH119322 to Cagri Yuksel.

## Author Contributions

X.C, F.D, and D.O designed the original studies the data was derived from.

X.C, L.W, E.C, F.D, and C.Y analyzed the data.

H.U.Y, X.C, I.G, and C.Y wrote the manuscript.

D.O and F.D provided feedback on the draft of the manuscript.

