## Supplementary Materials for "Poor Self-Reported Sleep is Associated with Prolonged White Matter T2 Relaxation in Psychotic Disorders"

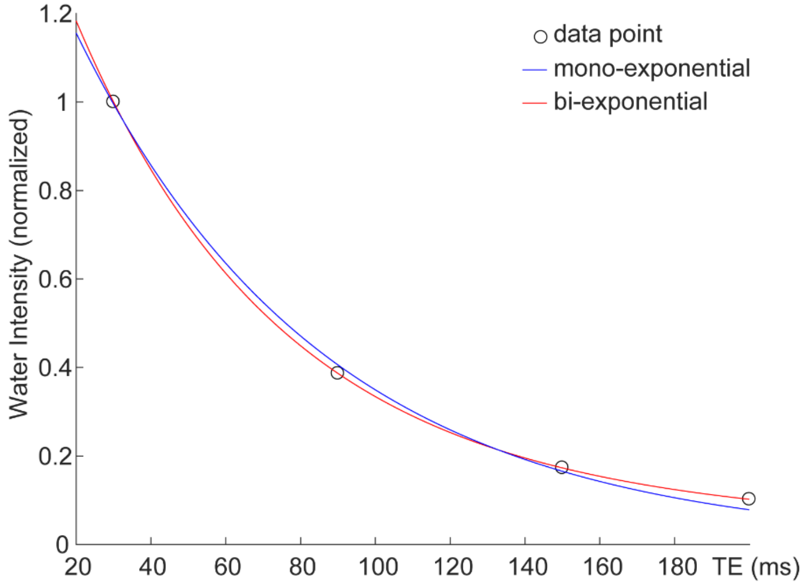


**Figure S1.** A representative T2 relaxation data set with mono-exponential and bi-exponential fittings

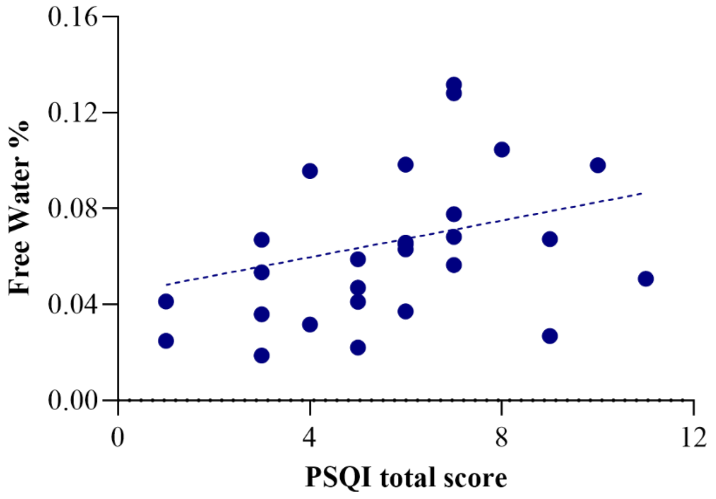


**Figure S2.** Correlation of PSQI total score with free water % (Spearman's rho=0.42).
